# Stochastic Simulation of Dopamine Neuromodulation for Implementation of Fluorescent Neurochemical Probes in the Striatal Extracellular Space

**DOI:** 10.1101/144436

**Authors:** Abraham G. Beyene, Ian R. McFarlane, Rebecca L. Pinals, Markita P. Landry

## Abstract

Imaging the dynamic behavior of neuromodulatory neurotransmitters in the extracelluar space arising from individual quantal releases would constitute a major advance in neurochemical imaging. Spatial and temporal resolution of these highly stochastic neuromodulatory events requires concurrent advances in the chemical development of optical nanosensors selective for neuromodulators in concert with advances in imaging methodologies to capture millisecond neurotransmitter release. Herein, we develop and implement a stochastic model to describe dopamine dynamics in the extracellular space (ECS) of the brain dorsal striatum. Our model is developed from first principles and simulates release, diffusion, and reuptake of dopamine in a 3D simulation volume of striatal tissue. We find that *in vivo* imaging of neuromodulation requires simultaneous optimization of dopamine nanosensor reversibility and sensitivity: dopamine imaging in the striatum or nucleus accumbens requires nanosensors with an optimal dopamine dissociation constant (*K_d_*) of 1 μM, whereas *K_d_* above 10 μM are required for dopamine imaging in the prefrontal cortex. Furthermore, our model reveals that imaging frame rates of 20 Hz are optimal for imaging temporally-resolved dopamine release events based on the probabilistic nature of dopaminergic terminal activity in the striatum. Our work provides a modeling platform to probe how complex neuromodulatory processes can be studied with fluorescent nanosensors and enables direct evaluation of nanosensor chemistry and imaging hardware parameters. Our stochastic model is generic for evaluating fluorescent neurotransmission probes, and is broadly applicable to the design of other neurotransmitter fluorophores and their optimization for implementation *in vivo*.

## INTRODUCTION

Monoamines such as dopamine, norepinephrine, and serotonin belong to a group of signaling molecules in the brain collectively known as neuromodulators. While classical neurotransmission is confined to communication between the pre- and postsynaptic neuron, and is mediated by fast acting ligand-gated ion channels, neuromodulation employs slower acting metabotropic receptors that exhibit a high level of extrasynaptic expression.^1^ Thus, modulatory neurotransmitter activity extends well beyond the synapse. As a consequence, neuromodulators such as dopamine influence a population of neurons beyond the synapse, enabling a single neuron to modulate the activity of a larger network of connections. It is therefore of great interest to develop tools to observe and quantify the release, diffusion, and reuptake of neuromodulatory neurotransmitters such as dopamine, where the spatial and temporal dynamics observed in the brain extracellular space (ECS) can be directly linked to receptor activation, neuronal activity, and behavior.

Among the most prominent dopaminergic systems are the nigrostriatal, mesocortical, and mesolimbic projections. Small clusters of dopamine neuron cell bodies located in the substantia nigra pars compacta (SNc) make extensive connections with the medium spiny neurons (MSN) of the dorsal striatum, forming the nigrostriatal pathway.^2^ This pathway is responsible for controlling fine motor movements and its dysfunction underlies the pathology of Parkinson’s Disease.^3^ Dopaminergic cell bodies in the ventral tegmental area project into the nucleus accumbens and the prefrontal cortex, forming the mesolimbic and mesocortical pathways, respectively.^2^ These systems play significant roles in cognitive control of behavior and reward processing, and their dysfunction underpins the pathology of depression, addiction, schizophrenia and attention deficit hyperactivity disorder (ADHD), among others.^4,5,6,7,8^ In all of these systems, neuromodulation, as opposed to neurotransmission, is the primary mode of influence. This diffusion-mediated transport of dopamine in the ECS is also known as volume transmission.^9^

One of the most ambitious pursuits in neuroscience is elucidating the relationship between neurons, neural circuits, behavior, and disease.^10^ Successful chronic and real-time recording of neurotransmitter mediated chemical signaling would be a decisive step in that direction.^11^ Current methods to measure the dynamics of dopamine volume transmission in ECS lack the spatial and/or temporal resolution of relevance to study neuromodulation. Voltammetry and amperometry are electrode-based methods used to record the presence of neuromodulators via redox chemistries, yet require penetration of the brain tissue and only assay neurotransmitter concentration at one point in space. Optical probes include cell-based neurotransmitter fluorescent-engineered reporters (CNiFERs) that have been engineered to express a chimeric dopamine receptor and a genetically encoded calcium indicator.^12^ CNiFERs utilize slow G-protein coupled receptor (GPCR) responses and thus do not report millisecond or micron-scale neurotransmitter activity. Fluorescent false neurotransmitters (FFNs) fluorescently label dopaminergic vesicles and provide single release site resolution but do not report neurotransmitter concentrations in ECS.^13-15^ Calcium imaging can show bouton activity in dopamine axons preceding release but tell us little about extracellular dopamine concentration.^15^ In sum, existing methods are insufficient to enable reliable measurements of dopamine and other modulatory neurotransmitters in the ECS with the necessary spatial and temporal resolution to be commensurate with physiological function. As a result, the theoretical framework developed to date has served primarily to describe the local, not global, dynamics of neurotransmission. Recent advances in new optical probes that report changes in extracellular dopamine concentrations in real time with significant spatial information warrant a thorough theoretical study of how such optical reporters of ECS neurotransmitter concentrations should be designed and implemented *in vivo* for recording fast dynamic processes.

Herein, we generate a theoretical framework with which to evaluate fluorescent probes designed to record the dynamics of modulatory neurotransmitters in the ECS under *ex vivo* and *in vivo* imaging conditions. We develop a model to probe the spatiotemporal profiles of dopamine in the striatum accounting for the stochasticity of dopamine quantal release, diffusivity, and reuptake, combined with fluorescent nanosensor kinetics and microscopic imaging parameters. We implement our model in the context of nanoparticle-based near-infrared fluorescent nanosensors for dopamine^16,17,18^ and develop appropriate neurochemical and imaging boundary conditions under which imaging of fast dynamics of dopamine neuromodulation can be accomplished for *in vivo* relevant spatial and temporal scales. Our results show that the process of dopaminergic neuromodulation occurs on characteristic timescales on the same order as exposure times used to optimize fluorescence imaging, thereby introducing temporal distortions in dopamine recordings. Therefore, when we optimize sensor-analyte binding interactions, a phenomenon emerges in which only sensors with kinetics parameters in a small critical window become feasible for *in vivo* imaging of fast dynamic processes. We probe these phenomena in the spatial domain as well. With appropriate *in vivo* firing behavior of striatal dopamine neurons, we show that optimally selected sensor kinetics and imaging frame rates can capture behavior-relevant dynamics of phasic firing in the dorsal striatum, with the necessary temporal resolution and signal-to-noise ratio to distinguish individual dopamine transient events elicited by release and reuptake.

## RESULTS AND DISCUSSION

We model the diffusion driven dynamics of dopamine in the ECS of striatal tissue by numerically solving the diffusion equation with dopamine source and sink terms. Dopamine sources are the quantal releases from dopaminergic terminals within a defined simulation volume, while the reuptake of dopamine from the ECS acts as a sink. Reuptake parameters are assumed to be uniform throughout the simulation volume. We set the simulation volume as a cubic block of striatal tissue comprised of uniformly distributed dopamine terminals. A schematic of striatal tissue model with dopamine terminals depicted as yellow spheres is shown in Figure 1a. We use the nigrostriatal projection as a model system owing to its critical role in reward and reinforcement, addictive behavior, habit formation, and its implications for motor neuron disorders, such as Parkinson’s disease. Furthermore, the nigrostriatal system is well studied in the literature, providing requisite physiological parameters relevant to dopaminergic neurotransmission with which to implement our model.

**Figure 1.**
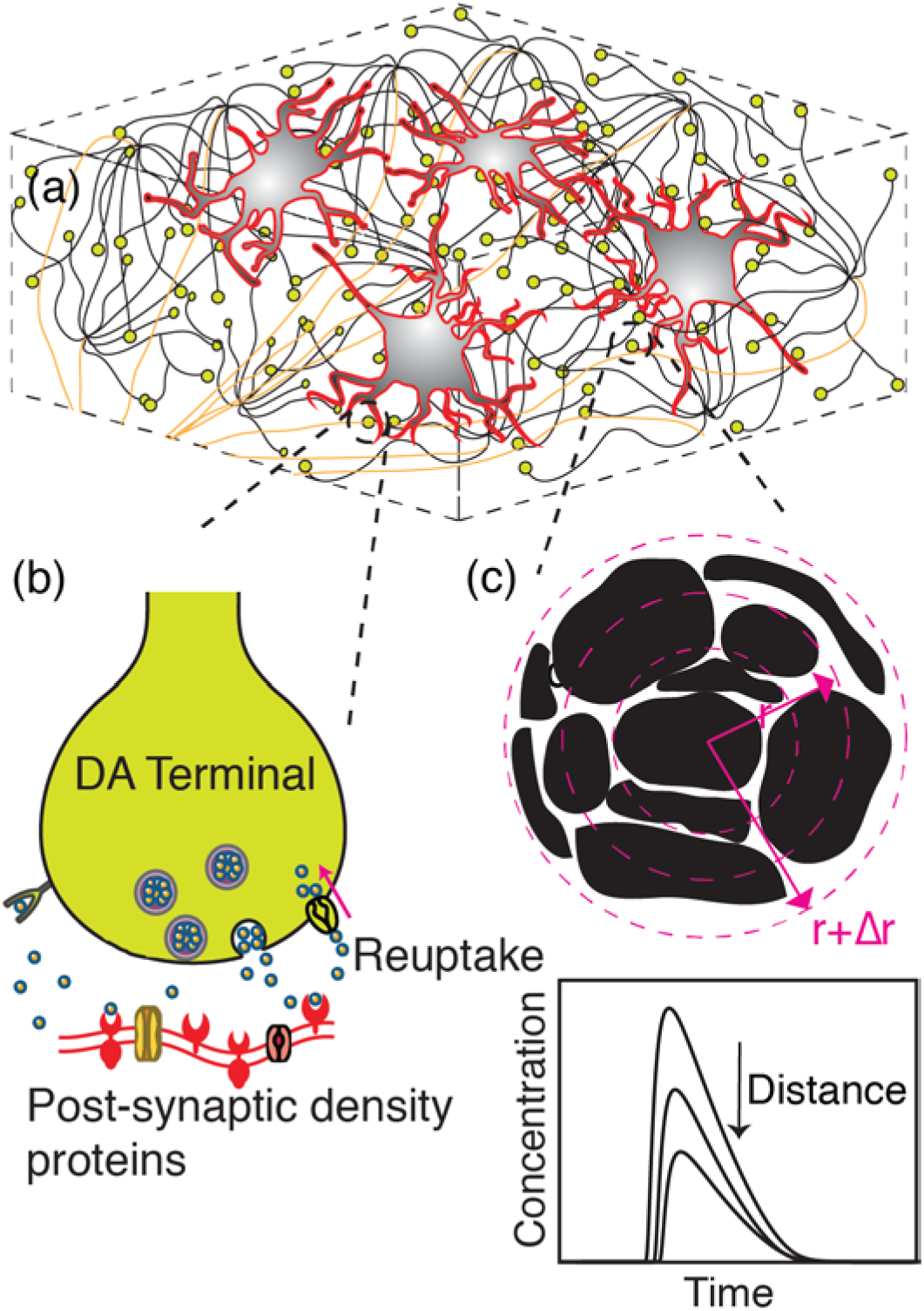
Schematic of dopamine model. **(a)** Dorsal striatum with medium spiny neurons (MSN - red contours/gray body), dopamine terminals (yellow), and projection axons (tan) from SNc **(b)** A magnified view of an individual dopamine terminal forming a synapse onto a dendritic shaft of MSN. Dopamine release: An action potential causes a dopamine-containing vesicle to release dopamine into the synaptic cleft. Dopamine encounters post-synaptic receptor proteins, triggering further downstream neuronal processes. Dopamine reuptake: DATs clear dopamine from the ECS to be recycled. **(c)** Space discretization around a dopaminergic terminal representing tortuous morphology of brain tissue. Black represents tissue surrounded by void ECS. Concentric circles depict simulation volume elements. Inset graph: Dopamine concentration fluctuates in space and time as a result of release, diffusion and reuptake.

### Simulation Volume

Dopamine terminals are the source of dopamine in our simulation, and dopamine transporters (DATs) drive dopamine reuptake. To elucidate the spatiotemporal dynamics of dopamine concentration in the ECS, we define a simulation volume as a ~10 μm^3^ section of striatal neural tissue containing 100 dopaminergic terminals. Terminals are arranged in a periodic lattice structure filling the simulation volume. The structural and functional parameters of our simulation volume are summarized thus: (i) the density of dopaminergic terminals in the striatum, (ii) probability of dopamine release upon membrane depolarization, (iii) amount of dopamine released per quanta (per vesicle fusion), (iv) effective diffusivity of dopamine in tissue and (v) dopamine reuptake kinetics by DATs. A summary of parameter values and literature sources is provided in Table 1.

**Table 1.**
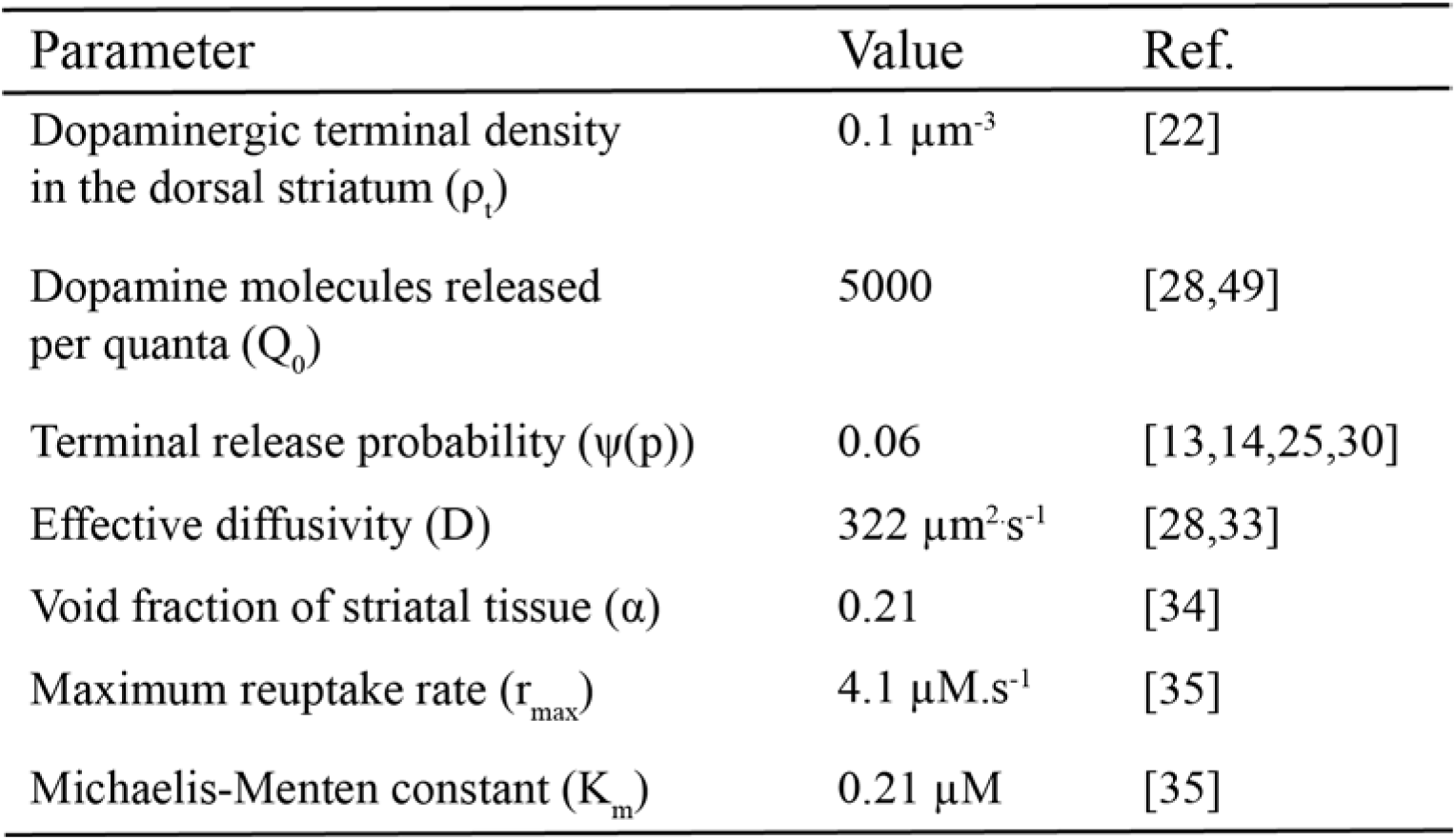
Values of Simulation Parameters and Literature Sources

### Model Representation of Dopamine Terminals

We define dopamine terminals as the boutons of axonal projections from the SNc into the dorsal striatum (Figure 1a). Cell bodies of dopaminergic neurons of the nigrostriatal pathway are located in the SNc and their axonal projections ascend into the dorsal striatum and make connections with dendritic spines or dendritic shafts of MSNs.^19-21^ These ascending axonal tracts are notable for their high terminal density, where each axon is estimated to make on the order of 400,000 connections in the striatum.^22,23,24^ A dopaminergic terminal contains a cluster of vesicles in close proximity to symmetric membrane densities, forming dopaminergic synapses of MSNs (Figure 1b). Our simulation considers these sites as point release sources in a three dimensional space. We recreate the neuroanatomy within the simulation volume as described in previous computational studies.^25-28^

### Dopamine Release Sites and Probability of Release

Neurotransmitter release occurs at release sites within synapses. Central synapses of the nervous system such as those found in the striatum contain a single release site, as comprehensively reviewed in Stevens et al.^29^. During an action potential, a single quantal release of dopamine occurs with a certain probability *p*, at each terminal (Figure 1b). Dreyer et al. calculate probability of 6% based on studies of neurotransmitter release using FFNs^13,14^ and the dopamine content of striatal tissue.^30^ The low dopamine release probability is consistent with experimental results, which revealed many dopamine terminals remain “silent” during stimulation.^13,14^ We assume a constant probability of release and quantity of release in the simulation, although some temporal and spatial heterogeneities have been reported.^13,31,32^ Thus, we execute our model for which each dopamine terminal possesses a single release site, where dopamine release probability per action potential per terminal is set to 6%^25^ and is assumed constant. Furthermore, membrane depolarization driving neurotransmitter release is mitigated by sodium pump activity, where sodium pumps remain inactive for ~ 10 ms following an action potential.^29^ Thus, we impose a constraint in our simulation to limit sequential dopamine release events to occur at intervals greater than 10 ms per terminal. As such, we impose a condition for the maximum release rate from a given terminal to 100 Hz. Despite this maximum release rate, the low probability of release makes it such that the 100 Hz boundary condition is rarely encountered in our simulations.

### Simulation of Release, Diffusion and Reuptake

Our simulation of dopamine concentration in the ECS invokes the equation of change for species conservation surrounding a dopaminergic terminal^27^

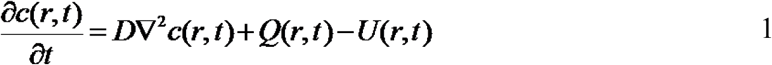

where *c(r, t)* represents spatial (*r*) and temporal (*t*) variation in dopamine concentration, and *Q(r, t)* and *U(r, t)* represent quantal release of dopamine into the ECS and reuptake by DATs, respectively. D is the effective diffusivity of dopamine in tissue after accounting for tortuosity of brain tissue.^33,34^ We solve this governing equation at each dopamine release terminal using finite difference method, and obtain the solution for temporal and spatial dopamine dynamics resulting from release from one dopamine terminal. Striatal tissue is composed of approximately 1 terminal per 10 μm^3^. *^22 24^* With this terminal density, we can determine the temporal profile of dopamine concentration resulting from the activity of all terminals included the simulation volume of interest. To do so, we solve the governing equation individually for each dopamine terminal, and superimpose temporal solutions of the governing equation to determine dopamine dynamics at any location within the simulation volume. Thus, the temporal change in dopamine concentration at any point in the ECS is the sum of the dopamine dynamics contributed by all terminals in the point of interest vicinity. We note that the error introduced by summing the non-linear reuptake term is negligible: dopamine re-uptake approximates linear behavior at sites distant from a release point, where the spatial summation occurs. Lastly, we discretize the governing equation to solve it numerically, since no known analytical solutions exist for this equation. The difference equation is written in spherical coordinates as forward difference in time and central difference in space. We use radial steps, *dr*, of 0.2 μm and time steps, *dt*, of 0.02 ms, which yield stable solutions over a wide range of biological parameters.

### Quantal Release

In equation 1, *Q(r,t)* represents quantal dopamine release following vesicle fusion^25^ and is represented by:

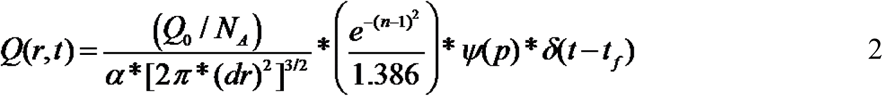

where *Q_0_* represents the number of dopamine molecules released per exocytosis event and *N_A_* represents Avogadro’s number. The parameter *ψ* assumes a value of 1 or 0 based on a release probability *p*. A release event increases the concentration of the first spatial element of the simulation volume (Figure 1c) by an amount represented by:

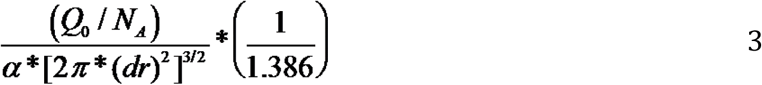

This is the volume element immediately surrounding the location of the dopamine terminal (Figure 1c). The parameter ∝ accounts for porosity of brain tissue.^27^ We use a Gaussian probability density function to determine the spatial distribution of dopamine immediately after release, normalized to ensure that only 5000 (*Q_0_*) molecules of dopamine are released per quanta (Table 1). Dopamine spillover after quantal release is instantaneous.^28^ Thus, a quantal release event affects the concentration of volume elements away from the center of the release site by an amount equal to the increase in the center of the release site (equation 3) scaled by an exponential decay term, 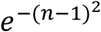. This exponential term is a function of the distance of the volume element from the center of the release site *r* = *(n-1)dr*, where *n* represents the spatial index, *n* = *1, 2*…*N* representing each volume element in the simulation. Firing frequency (F) sets the number of action potentials over a given simulation period. The temporal distribution of action potentials over the simulation time period, *t_f_*, is modeled as a Poisson distribution with mean a firing rate of *F*. *δ*(*t* − *t_f_*) is a delta function in time and ensures that release can only occur during an action potential firing event with a binary probability *ψ*.

### Dopamine Reuptake (U)

Uptake of dopamine from the ECS occurs via DATs. In our model, we assume a uniform distribution of DATs in the simulation volume and model dopamine uptake with a Michaelis-Menten rate equation with parameters *r_max_* and *K_m_* and in a medium of porosity *α*.^27,35^

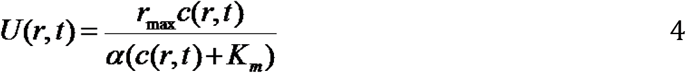

Dopamine saturation must be taken into account especially for simulation regions in close proximity to a terminal, where dopamine concentration immediately following a release can be very high. This non-linear uptake expression (equation 4) allows for saturation of the dopamine reuptake process at high physiological dopamine concentrations, corresponding to the case of substrate saturation.

### Single Terminal Behavior

The spatiotemporal dynamics of dopamine in the ECS following release from a single terminal influences dopamine receptors within the diffusion volume prior to dopamine reuptake.^28^ Dopamine is the primary endogenous ligand for two dopamine receptor sub-classes: D_1_-type and D_2_-type receptors, with EC_50_ binding affinities of 1 μM and 10 nM respectively.^28,36^ We define the sphere of influence of a quantal release of dopamine after release from a terminal based on these activation EC_50_ values.^28^ Our simulation shows that for a single quantal release from a terminal, the sphere of influence on low affinity D_1_-type receptors and high affinity D_2_-type receptors is 7 μm and 17 μm, respectively (Figure 3a). The spatiotemporal dynamics of a quantal release from a single terminal over a 20 μm radial distance shown in Figures 2a,b, Figure 3a and Figure S1, is consistent with prior studies that show that dopamine diffuses from synaptic termini in quantities that overflow the synaptic cleft, giving rise to dopamine volume transmission.^26, 28^

Dopamine propagation from the center of the synaptic cleft occurs on short time scales relative to dopamine diffusive effects. Our simulation shows that D_2_ and D_1_-type receptors within the sphere of influence of a terminal are activated within 50 ms and 20 ms of dopamine release, respectively (Figure 2c). Furthermore, we compute the speed of propagation of receptor activation as a function of distance from the release site (Figure 2d). The D_2_-type receptor activation wave front (10 nM dopamine) moves forward at peak speeds approaching 1.5 μm/ms and slows to 0 μm/ms as a function of distance from the release site. Similarly, we observe peak propagation speeds of 1 μm/ms for D_1_-type receptor activation (1 μM of dopamine). Our simulations provide estimates for speeds of dopamine volume transmission, and the time-dependence of dopamine’s activity on distal receptors from the release point. Surprisingly, dopamine volume propagation speeds we compute for D_1_- and D_2_-type receptors is four to five orders of magnitude slower than reported speeds of electrical signal propagation in nerve fibers.^37,38^ To our knowledge, this is the first quantitative report on the speed of chemical signaling of dopamine in the dorsal striatum.

**Figure 2.**
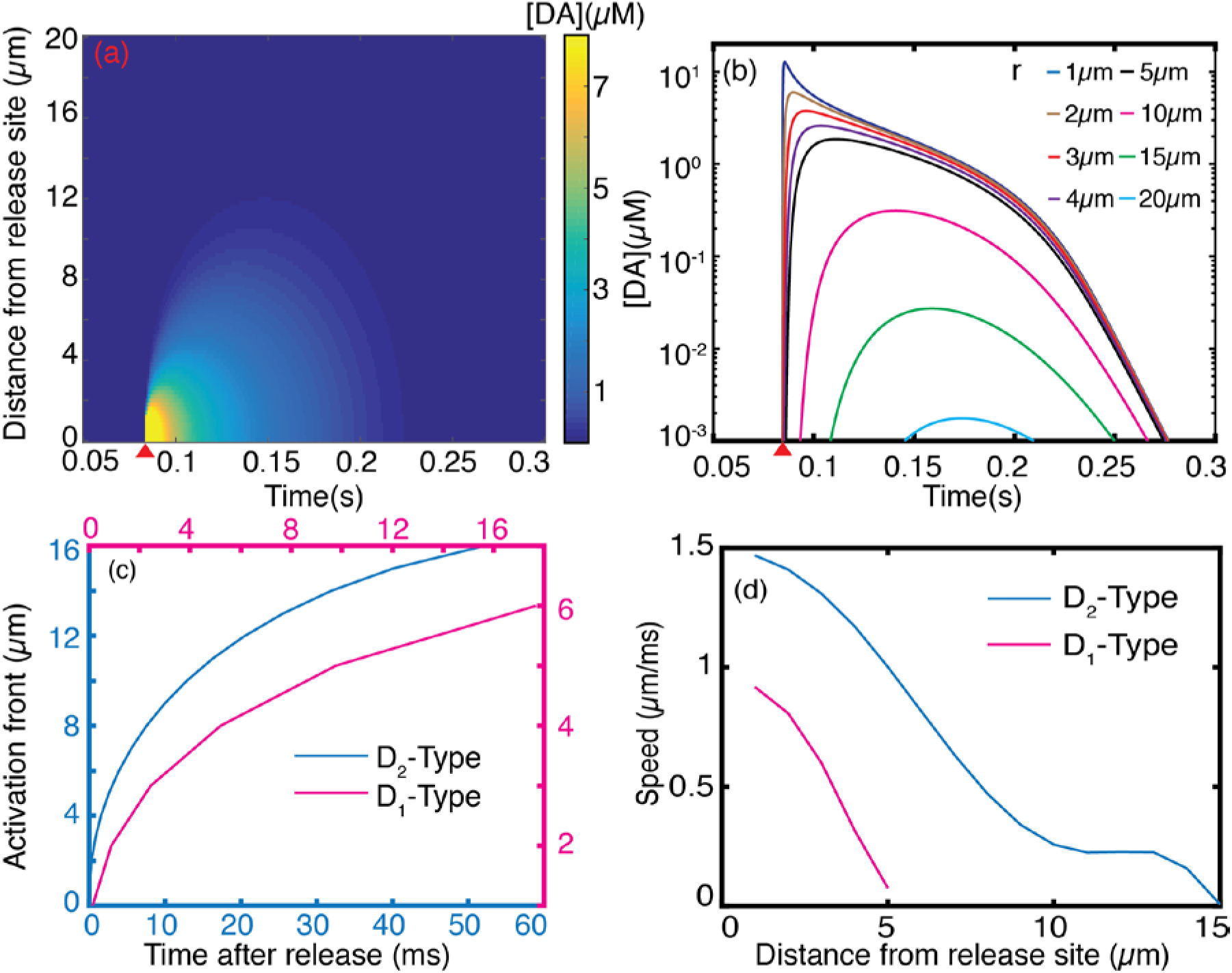
Spatiotemporal dopamine dynamics following a single action potential driven quantal release of dopamine. (a) Dopamine concentration profile evolution following a single quantal release as a function of distance and time. Red wedge indicates quantal dopamine release. (b) Dopamine spatial concentration profile at varying distances from release site. (c) Front (instance where EC50 is exceeded) of dopamine receptor activation following dopamine release for D1 (pink) and D2 (blue) type receptors. (d) Propagation speed of D1 and D2-type receptor activation after a quantal release obtained from first derivative of (c).

Non-linear reuptake kinetics we defined in Equation 4 is sometimes approximated by a linear approximation of the form (*r_max_*/*K_m_)c(r, t)*.^27^ While a linear approximation of dopamine reuptake facilitates an analytical solution for the governing equation, linearization creates significant deviation from dopamine dynamic behavior obtained with non-linear reuptake kinetics (Figure 3a). The impact of the linear approximation on dopamine reuptake kinetics arises from neglecting the fact that DATs in close proximity to the releasing terminal are saturated (*c(r, t)* >> *K_m_*) and can only clear dopamine at a maximum rate of *r_max._*. The linear approximation overestimates uptake in regions proximal to the point of dopamine release, resulting in different linear versus non-linear reuptake kinetics. Our model implements non-linear reuptake kinetics with a Michaelis-Menten rate equation, and enables us to calculate the spatial sphere of influence with the biologically-relevant influence of saturating dopamine reuptake proximal to the release site. Indeed, our results show the spatial sphere of influence with linear uptake is only half of that obtained with non-linear kinetics for D_1_-type receptors (Figure 3a). To demonstrate the importance of treating the reuptake kinetics as a non-linear saturable process, we compared the linear model at nominal *r_max_* and *K_m_* values (Table 1) with a non-linear model in which *r_max_* was increased by a factor of 10 (equivalent to increasing the density of DATs by an order of magnitude) and found the two dynamic behaviors to be comparable (Figure S2). Our results exemplify the importance of treating dopamine clearance from ECS as a non-linear process, especially when spatial and temporal domains are simultaneously considered, and provide a quantitative comparison between linear and non-linear reuptake on the dynamics of dopamine in ECS.

**Figure 3.**
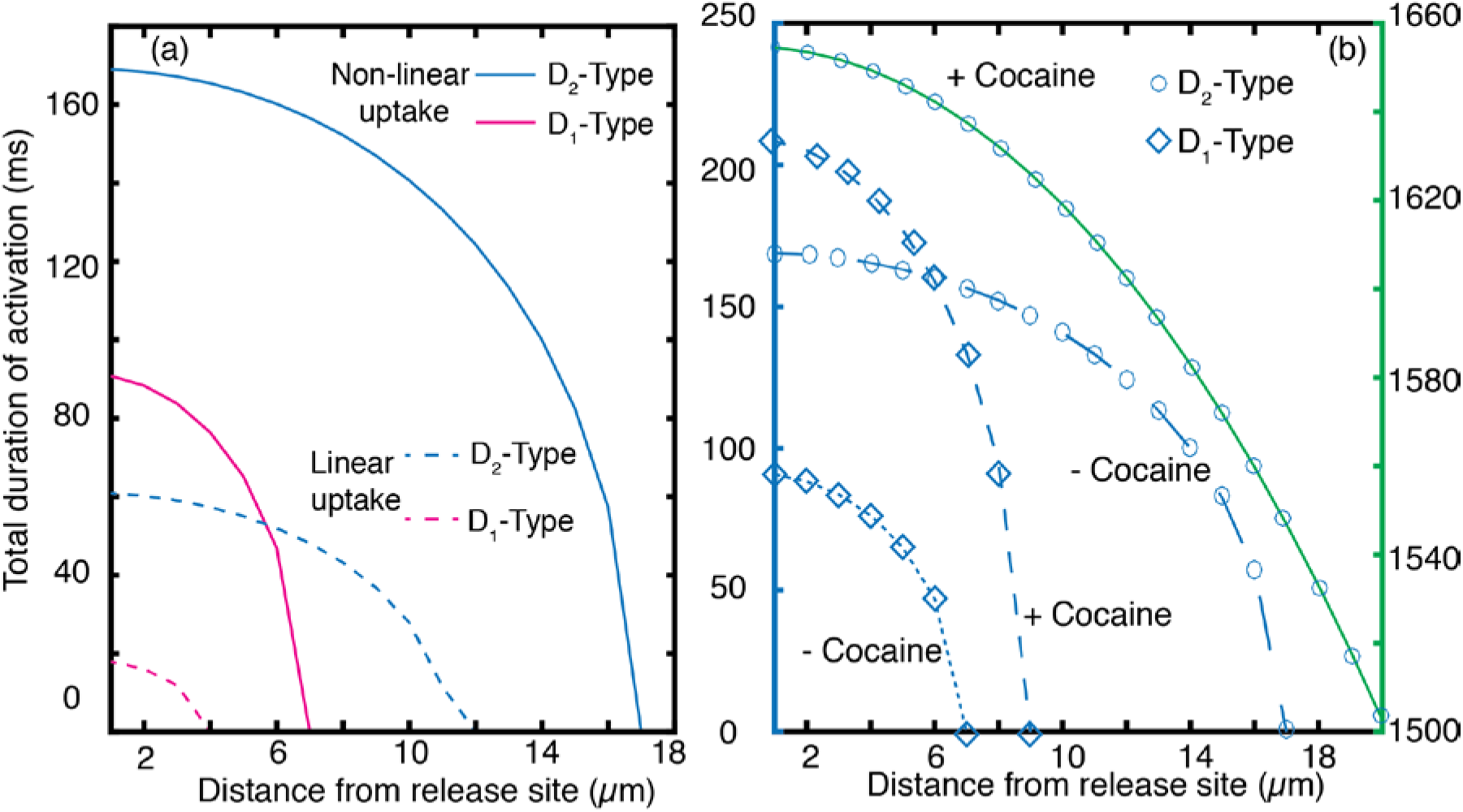
Non-linear computation of receptor activation by dopamine diffusion. (a) Sphere of influence of a single dopaminergic terminal over D_1_- and D_2_-type receptors. Time over which the EC50 of each receptor type is exceeded is plotted as a function of distance from the terminal. D_1_- type receptors are insensitive to quantal release at 7 μm radial distance beyond the release site, whereas D_2_-type receptors can be influenced by a single quantal release through a radial distance of 17 μm. The duration during which the EC_50_ of each receptor type is exceeded decreases monotonically with distance. Broken lines represent spheres of influence of linear uptake kinetics with the nominal *r_max_* and *K_m_* values listed in Table 1. (b) Effect of dopamine reuptake blocker, cocaine, on sphere of influence, plotted as receptor activation duration as a function of distance from release site. Competitive inhibition by cocaine increases the sphere of influence of receptors dramatically, where the green scale on the right y-axis shows values for cocaine-affected D_2_-type receptors.

The mechanism of action of many therapeutic drugs and most drugs of abuse are similar: drugs competitively bind to DATs and modulate dopamine clearance rate from the ECS. We demonstrate the applicability of our model to probe the neurophysiology of dopaminergic systems by probing the effect of dopamine reuptake inhibition by cocaine. Cocaine has been shown to increase the affinity parameter in equation 4, *K_m_*, from 0.21 μM to 8 μM.^39^ Our simulation shows that inhibition of dopamine reuptake by cocaine increases the duration of D_1_-type receptor activation by up to two fold and that of D_2_-type receptors by up to six fold (Figure 3b). The spatial spheres of influence from a quantal release increase from 7 μm to 9 μm, and from 17 μm to 47 μm, for D_1_ and D_2_-type receptors, respectively. Our results suggest few competitive inhibitors – whether therapeutic or abusive – have drastic effects on the dopamine sphere of influence for a singular dopamine terminal, achievable when we model dopamine reuptake as a non-linear process.

### Many-Terminal Behavior

Dynamic dopamine behavior at a point in the striatal ECS is influenced by the behavior of all active terminals in the vicinity. Dopaminergic neurons exhibit slow tonic and fast burst firing activity.^40,41^ A burst in firing activity correlates with reward reinforcement as a response to salient events, whereby striatal dopamine neurons burst in spike trains of 4 to 7 spikes per burst event at a spiking frequency of 20 Hz.^40,42,43^ Conversely, a pause in firing is correlated with response to adverse events or withdrawal of an expected reward. Tonic activity underlies dopaminergic activity at rest.^40,41,42,43^ We implement our model to calculate dopamine concentrations in a volume of striatal brain tissue for a simulated spike train of physiological relevance. To account for the neurologically relevant case of collective multi-terminal activity, we extend our model to employ spatiotemporal summation of solutions from each terminal surrounding of a point of interest in the ECS. To this end, we evaluate spatiotemporal dopamine dynamics at a point of interest surrounded by 100 dopamine terminals (Figure 4a) and, separately, at a point surrounded by 25 terminals (Figure 4b) arranged at uniformly spaced cubic lattice points, with no terminal located closer than 2 μm to the point of interest. The terminal spacing of each cluster is based on density parameter defined in Table 1. We chose an extrasynaptic point located at least 2 μm from the closest terminal to avoid capturing the dominant behavior of synaptic dopamine hot spots in which behavior is dominated by the firing activity of the closest terminal. We simulate a 2-second spike train representative of phasic firing behavior by implementing our model over four distinct firing regimes (Figure 4a, b). The simulated firing frequency and duration is chosen based on experimentally observed *in vivo* spiking activity of dopaminergic neurons^40,42^: we simulate an ensemble of dopaminergic neurons undergoing a 4 Hz tonic firing rate for t = 0 to t = 0.4 s, followed by a 20 Hz burst firing regime for t= 0.4 s to t = 0.7 s, followed by a 0.5 s pause (0 Hz) until t = 1.2 s. For the last 0.8 s of the simulation, we return to a 4 Hz tonic firing regime. Note that each firing rate is the mean of a Poisson distribution of firing rates among the ensemble, as we described in preceding sections. The simulation ensemble size is commensurate with experimental fast scan cyclic voltammetry (FSCV) assays where the carbon fiber electrode samples dopamine overflow from a region encompassing ~100 dopaminergic terminals.^44,45^

**Figure 4.**
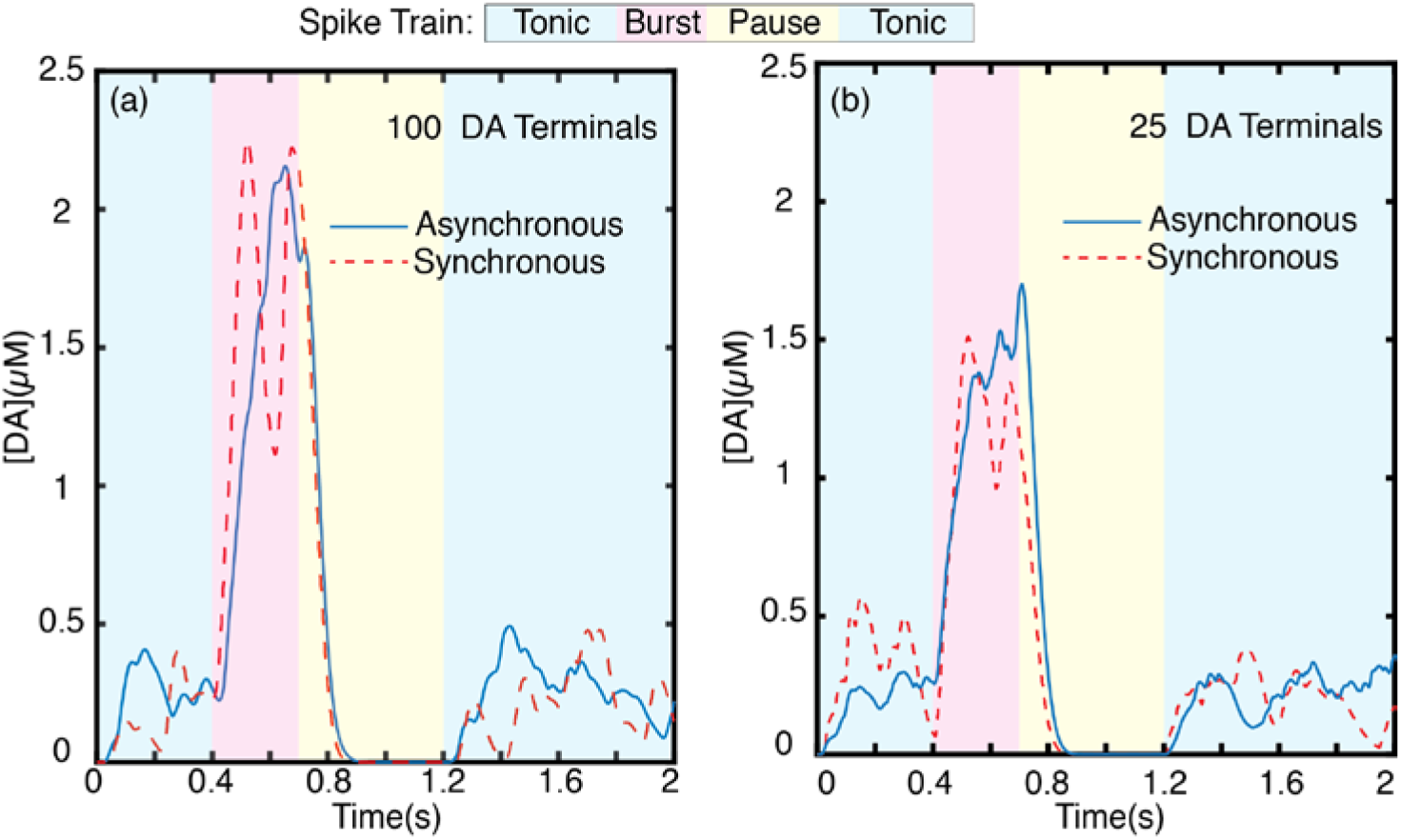
Dopamine concentration evolution profile for a simulated volume. (a) Dopamine dynamics at an extrasynaptic point surrounded by (a) 100 and (b) 25 phasically firing dopaminergic neurons with asynchronous firing (blue trace, solid) and synchronous firing (red trace, dash). Each trace represents the average of N=20 independent simulation runs.

We present results from a cluster of 100 dopaminergic terminals and 25 dopaminergic terminals firing phasically with the above-described spike train to highlight how the underlying functional connectivity of terminals can result in different spatiotemporal dopamine behavior. While the firing activity of an individual or a pair of dopaminergic neurons is well studied, the number of terminals in a phasically firing ensemble is not well understood.^42,46^ We predict that functional or sub-functional connectivity influences the size of phasically firing ensembles and their synchrony.^42^ As such, we simulate a spike train from a phasically firing cluster of 100 dopaminergic terminals (Figure 4a) compared to a cluster of 25 dopaminergic terminals (Figure 4b) firing synchronously or asynchronously. Dopamine release is a highly stochastic process and the results presented here are average behavior from N = 20 separate runs of our simulation. We present individual simulation runs and run average for the 100 terminal asynchronous firing case in Figure S3. In both terminal clusters, asynchronous firing results in a more temporally homogenous concentration profile. Synchronous firing concentration profiles exhibit sharp dopamine concentration transience in all firing regimes (Figure 4a, b). Peak dopamine concentration during burst phase is modestly higher for the 100 terminal cluster, and its scale and diffusion also has a larger spatial extent. Tonic dopamine concentration levels, however, are roughly the same for both the 100 and 25 terminal clusters. For both simulated terminal cluster sizes, tonic asynchronous firing gives rise to a steady basal dopamine level whereas synchronous firing does not. The pause in firing following burst firing activity clears dopamine from the ECS in both cases; complete clearance is achieved within 150 ms of the onset of pause in firing. The observation that tonic dopamine concentrations are mediated by uncorrelated, asynchronous firing is in agreement with prior studies, which show that tonic activity gives rise to the basal dopamine level measured in ECS.^8^ It is worth noting the concentration profile depicted in Figure 4 measures dopamine for a singular point in ECS. Of relevance to the spatial and temporal limitations of existing experimental tools to probe brain neurotransmission, we implement our model for space averaged dopamine dynamics, as detailed below.

### Dopamine Nanosensors in the Striatal ECS

Of relevance to fluorescent probes for measuring neurotransmitter concentrations in the ECS,^16^ our model captures spatial evolution of dopamine in the ECS, in addition to the temporally-relevant information obtained from FSCV. Nanoparticle based optical probes hold great promise for probing volume transmission dynamics in the ECS in a space and time resolved manner. First, nanoparticles can fit into the intricate porous and tortuous morphology of the ECS, allowing them proximate access to synaptic and extrasynaptic locations to record dynamic concentration behavior. Second, their diffusive distribution over large volumes of the ECS provides much needed spatial information. Here, we develop a model of the sensor’s fluorescence modulation in response to dynamic analyte behavior. In particular, we evaluate the dynamic range of a carbon nanotube-based dopamine nanosensor, previously validated for dopamine responsivity *in vitro*.16,47 Briefly, the polymer-nanotube conjugate nanosensor contains a surface environment with active recognition sites for dopamine.^16,48^ The reversible adsorption of individual dopamine molecules onto the nanosensor recognition sites increases the quantum efficiency of the nanotubes, causing a brightening and thus providing a fluorescent recognition signal. Denoting the total number of dopamine binding sites as L, the free (dopamine unbound) sites as *, the dopamine-bound sites as DA* and free dopamine molecules as DA, we establish equilibrium conditions for reversible dopamine adsorption to an ensemble of nanosensor binding sites in the ECS:

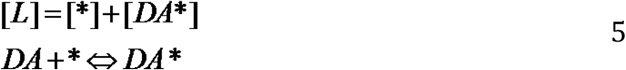

The first expression represents the dopamine active site balance and the second approximates that the dopamine adsorption process equilibrates on relatively short time scales compared to dopamine diffusion timescales in tissue. We define equilibrium constant, *Keq* for the dopamine-sensor binding process as:
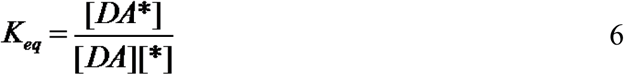

We next substitute the equilibrium constant into the site balance equation to derive an expression for the dopamine nanosensor fluorescent response as follows:
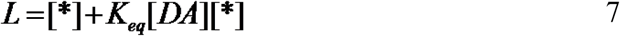

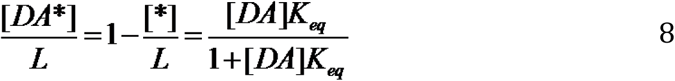

We note that the increase in nanosensor fluorescence intensity is directly proportional to bound active sites 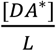. Thus, the expression for change in intensity normalized against initial nanosensor fluorescence 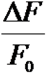 can be represented as:

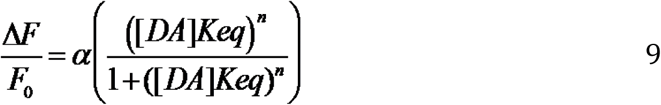

The additional fitting parameter, *n*, is introduced to account for dopamine binding cooperativity as determined for the nanosensor calibrated *in vitro* in Kruss et al.: =0.55, *n*=0.66, *Keq* = *2.31(μM)^-1^*.

We first implement our model to evaluate the spatial and temporal evolution of dopamine concentration following a dual quantal release from a terminal (Figure 5a). We probe dopamine concentrations at several distances from the terminal, (Figure 5 b,c) selected to be within the D_1_-type and D_2_-type receptor spheres of influence. Our simulation shows that the spatial dependence of dopamine concentration evolution and corresponding nanosensor response requires a 20 Hz frame rate to discriminate between two sequential release events occurring 0.2 seconds apart (Figure 5b, c).

**Figure 5.**
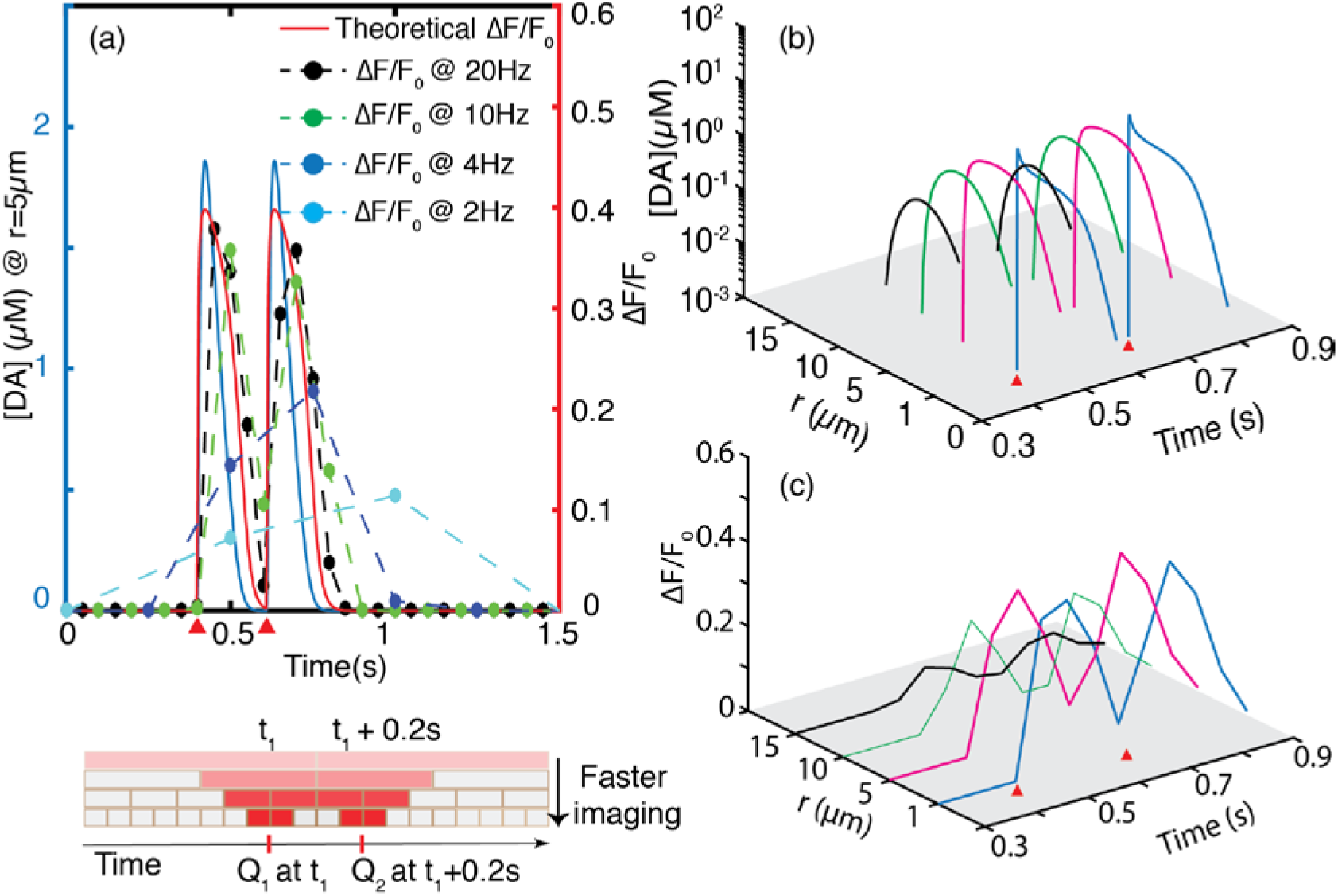
Temporal resolution of single and dual quantal dopamine release events. (a) Faster imaging frame rates enable resolution of quantal release events. As imaging frame rate increases, the observed nanosensor response more accurately captures the theoretical nanosensor response. Red wedge indicates time of quantal release. Bottom panel: schematic shows how faster imaging more precisely localizes temporal position of a single quantal release. Color gradient is to scale, showing the ΔF/F0 relative to the theoretically expected at each frame rate. (b) Discrimination between two quantal releases improves as one moves away from release site. Red wedges show times of quantal release positioned 0.2 s apart. (c) a 20 Hz frame rate resolves the two releases up to 15 μm away from the release site.

### Temporal Resolution Is Determined by Nanosensor Kinetics and Imaging Frame Rate

Nanosensor technologies to measure neurotransmitters in the ECS must capture hundred millisecond-scale dopamine release and clearance, as shown by our simulations. Temporally resolved neurotransmitter measurements with FSCV need only account for temporal sampling rates, which are easily achieved with a high scan rate voltammogram. Conversely, for nanosensors with fluorescence readouts, both temporal and spatial sampling rates will influence the measurement signal-to-noise, due to hardware limitations in fast sampling rates. Our above simulations set the physiologically relevant dopamine spatiotemporal dynamics in the striatum. Henceforth, we consider nanosensor performance limitations that are imposed by imaging hardware. While our model can be implemented generically for any fluorescence probe including calcium dyes, CNiFERs, and FFNs, among others, we focus on implementing our simulations for dopamine measurements in the striatal ECS using near-infrared fluorescent dopamine nanosensors.*^16^* During video-rate fluorescence imaging, substantive deviations from theoretical nanosensor response profiles are likely to be encountered owing to the short time scales over which dopamine is released into and cleared from the ECS. Specifically, quantal release and related dopamine dynamic behavior (Figure 5a, b, c) occurs on similar timescales as the exposure time used in conventional fluorescence microscopy (tens to hundreds of milliseconds). We must therefore account for the temporal distortion imposed on nanosensor response by imaging hardware. The nanosensor ΔF/F0 observed using a video-rate fluorescence imaging is evaluated as:
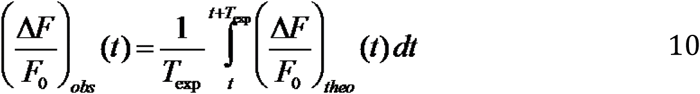

where 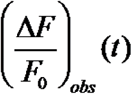 is the observed nanosensor response when imaging by fluorescence microscopy, and 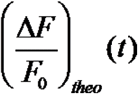 is the theoretical nanosensor response function. *T_exp_* is the camera exposure time and is inversely related to the nominal frame rate of imaging.

When camera exposure times are thus incorporated into our model, we indeed find that longer exposure times decrease the nanosensor fluorescence intensity recorded by the camera. As expected, the recorded spatial and temporal responsivity of the dopamine nanosensor underestimates the physiological dopamine concentration, and misses the true temporal release by overestimating the peak dopamine release time (Figure 5a, 5b). We compare the discrepancy between true nanosensor response and imaged nanosensor response for 2, 4, 10, and 20 Hz frame rates. The ability of a nanosensor to capture single terminal quantal release is compromised at frame rates below 2 Hz, where observed nanosensor response is only 20% of the predicted peak response, and could introduce a time-delay of up to 0.5 s. Conversely, when imaging with a 20 Hz frame rate (corresponding to a nominal 20 frame/s or 50 ms exposure time), 95% of the nanosensor fluorescence is imaged, and time-delay of no more than 50 ms is introduced between the quantal release event and the nanosensor response. Considering that there is diffusion induced temporal distortion of 30 ms at r = 5μm (Figure S1), this temporal delay at 20 Hz imaging becomes negligible. Of physiological relevance, quantal release events are expected to occur at least 0.2s apart in striatal dopaminergic terminals during tonic firing.^41,49^ When two quantal releases are located 0.2 s apart, a 20 Hz frame rate is needed to identify the two release events (Figure 5a). Conversely, both a 2 Hz and 4 Hz frame rate enables the nanosensor to record a spike in local dopamine concentration, but cannot discern that this spike is a result of two distinct quantal release events (Figure 5a). It is worth noting that the nanosensor intensity SNR is inversely proportional to the frame rate used to acquire the nanosensor signal, due to temporal averaging of signal photons. As a result, imaging frame rates cannot be increased infinitely to fully recapitulate the temporal profile of dopamine release. Below, we model optimization of spatial and temporal information in consideration of nanosensor SNR.

Our results show that diffusion of dopamine out of the synaptic cleft and into the brain ECS can be detected by nanosensors located up to 16 μm away from the terminal with ΔF/F0 of 5% or more when imaged at 20 Hz frame rates. Furthermore, we exemplify how nanosensors can be implemented to image temporal heterogeneities of dopamine dynamics, provided imaging hardware with sufficiently high frame rates. Specifically, frame rates of 20 Hz are needed to resolve single and multiple quantal releases that occur 0.2 s apart from the same terminal. In general, quantal releases located *x* ms apart require camera exposure times of less than *x/2* ms to be resolved. Slower frame rates improve SNR but decrease nanosensor response (ΔF/F0) and introduce significant temporal distortion on the measured dopamine response profile.

To optimize nanosensor performance, we tuned several parameters in the dopamine nanosensor model (equation 9). Our goal is to determine which nanosensor parameters will enable us to resolve dopamine dynamics of physiological relevance in the striatum. The equilibrium constant (*Keq*), the proportionality factor (α), and the cooperativity parameter (*n*), are intrinsic to the nanosensor and can be tuned to optimize nanosensor performance. The parameter α weighs the nanosensor quantum yield toward the nanosensor SNR. Higher α corresponds to stronger turn-on response (ΔF/F_0_) and improves SNR over the entire physiological dopamine concentration range. *K_eq_* is a measure of the affinity between the nanosensor and dopamine analyte. High *Ke*q (or low dissociation constant, *K_d_*) improves nanosensor response at low concentrations of dopamine, but also leads to quicker nanosensor saturation. On the other hand, low *Ke*q results in a nanosensor that is unresponsive to low concentrations of dopamine. Thus, to maximize the dynamic range of the dopamine nanosensor for the range of experimentally relevant dopamine concentrations of 30 nM – 10 μM^31,50,51^ we sought to identify nanosensor parameters that are amenable to capturing *in vivo* endogenous dopamine dynamics. First, we set a 5% ΔF/F0 lower-limit at the spatial boundary of the D_2_-type receptor sphere of influence (17μm), and this fixes *α* = *2*. We set the cooperativity factor n =1, which has been shown to represent our concentration range in experimental nanosensor calibrations*^16^* yielding a sigmoidal response curve representative of a Langmuir surface. Fixing *α* and *n*, we vary *K_eq_* over several orders of magnitude and determine *K_eq_* = *1 (μM)^-1^* to be optimal for a fluorescent nanosensor, which balances nanosensor reversibility and sensitivity according to biologically imposed boundary conditions. To evaluate nanosensor reversibility, we simulated nanosensor response to two quantal release events located 0.2 seconds apart from the same terminal (Figure 6a), imaged with a 20 Hz frame rate. We define reversibility as the fall in nanosensor intensity during clearance of the first quantal release, divided by rise in intensity in response to the first quantal release. The parameter *α = 2* sets maximum nanosensor response, and we define sensitivity as the measured peak ΔF/F0 divided by *α*. With these definitions, we varied *K_eq_* over several orders of magnitude to develop the parameter maps shown in Figure 6b and Figures 7a, b. High *K_eq_* values enhance sensitivity, enabling the nanosensor to turn-on at low dopamine concentrations. However, high *K_eq_* values cause nanosensor to saturate rapidly and adversely impact nanosensor reversibility (Figures 6a, b). On the other hand, low *K_eq_* values have very good reversibility but reduced sensitivity. At *K_eq_* = *1 (μM)^-1^*, we observe that the nanosensor both responds instantaneously to quantal dopamine release, and also captures dopamine reuptake kinetics to accurately discern between two quantal release events 0.2 seconds apart. Thus, we identify *K_eq_* = *1 (μM)^-1^* as optimal for imaging dopamine dynamics in the dorsal striatum, in which the fastest sequential quantal release events occur at least 0.2 seconds apart.^49^

**Figure 6.**
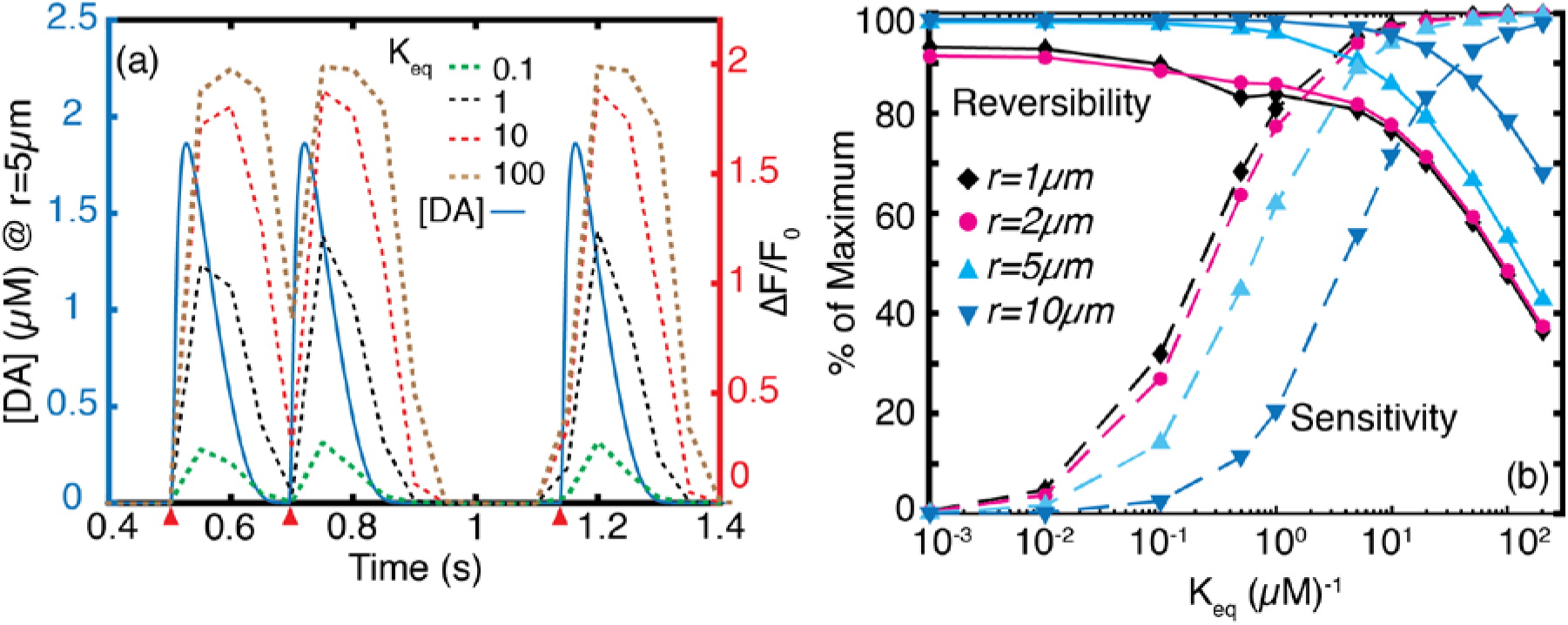
Effect of nanosensor parameter *Keq* on performance. (a) Dynamics of three quantal release events (red wedges) imaged using nanosensors for which *K_eq_* varies over three orders of magnitude. The first two quantal releases are located 0.2 s apart. At *K_eq_* = 100 *(μM)^-1^* the nanosensor affinity for dopamine is too strong, which adversely affects reversibility. The second release event cannot be resolved. Peak ΔF/F_0_ values increase with increasing *K_eq_*. At low *K_eq_* values, nanosensor shows high reversibility but poor sensitivity (b) Parameter space for reversibility and sensitivity at r = 1 μm, 2μm, 5μm and 10μm from release site. High dopamine concentrations proximal to the release site yield high percent nanosensor responsivity. However, maintaining nanosensor reversibility suffers proximal to the release site.

The parameter space we developed to optimize spatiotemporal signal acquisition of dopamine nanosensors with various *K_eq_* values can now allow us to test how camera frame rates affect the nanosensor reversibility and sensitivity parameter space. As we show in Figure 7a, fast imaging frame rates are needed if the nanosensor binds the dopamine analyte too strongly (large *K_eq_*) to temporally resolve the two quanta released 0.2 s apart. Therefore, we conclude that an optimal frame rate is a necessary but not sufficient condition to resolve the temporal heterogeneities of dopamine dynamics in the striatum. The nanosensor’s chemical responsivity and adsorption/desorption kinetics, combined with the imaging hardware limitations, both contribute to the spatiotemporal profiles of dopamine evolution that can be captured. When sequential release events faster than 0.2 s apart are considered, the reversibility curves shift towards nanosensors with lower *K_eq_* in a manner similar to that observed for imaging close to the release site (Figure 6b). Therefore, recording of faster dynamic events demands nanosensors with lower *K_eq_*, and comes with an opportunity cost of low sensitivity (Figure 6b) and higher noise (Figure 7c, Figure S4). The second chemical parameter of the nanosensor, the turn-on response parameter *α*, has little effect on nanosensor sensitivity and reversibility (Figure 7b), and which we set at *α* = 2.

**Figure 7.**
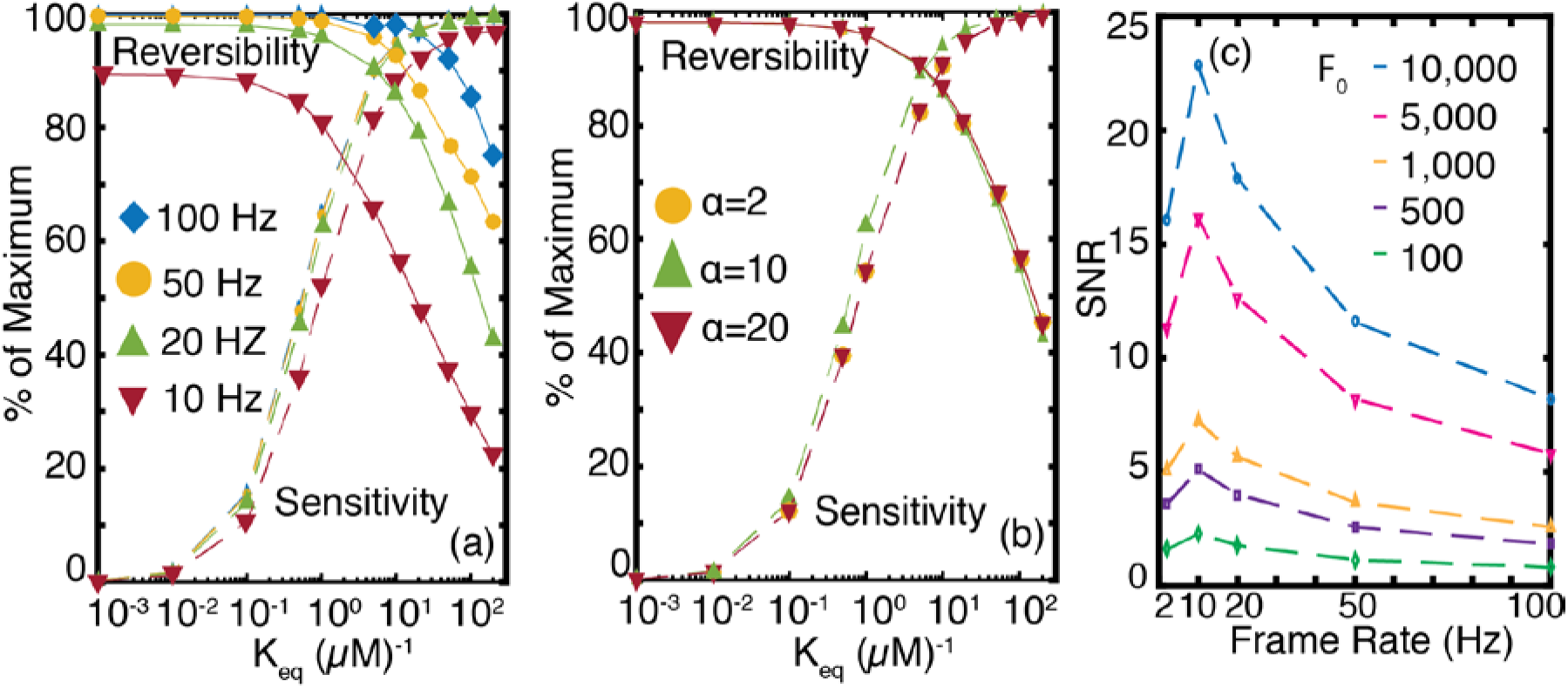
Nanosensor sensitivity, reversibility, and signal-to-noise ratio, probed for varying frame rates, nanosensor chemistries, and baseline nanosensor fluorescence. (a) Nanosensor parameter space for frame rates ranging from 10 Hz to 100 Hz with α = 2 (b) Nanosensor parameter space is largely insensitive to nanosensor turn-on response parameter *α*. (c) SNR shows strong dependence on frame rate (abscissa) and baseline intensity (F_0_, dashed traces) of the nanosensor. As frame rate increases, SNR passes through a maximum at 10 Hz and monotonously decreases afterwards. We note that 10 Hz maximizes SNR. The baseline fluorescence intensity F_0_ is varied from 100 to 10000 and corresponds to the 2 Hz frame rate, and units of F_0_ are arbitrary.

### Nanosensor Fluorescence and Imaging Frame Rate Considerations for Optimizing Signal-to-Noise Ratio

The observed nanosensor signal (ΔF/F_0_) takes into account the relationship between hardware (frame rate & instrument noise), and nanosensor chemistry (nanosensor fluorescent signal, sensitivity and reversibility parameters). During analysis of temporal resolution, we noted that frame rates could not be made arbitrarily faster because of their adverse impact on SNR. Here, we provide an analysis to include noise into the observed ΔF/F_0_. To elucidate how hardware and nanosensor chemistry contribute to SNR, we consider the peak signal from a nanosensor with optimal kinetic parameters of α = 2 and K_eq_ = 1(μM)^-1^ responding to a single quantal release of dopamine at a radial distance of r = 5 μm. The mathematical derivation of noise for ΔF/F_0_ is provided in Methods section. We consider the primary contribution of noise to the SNR to be Poisson noise (also known as shot noise) and do not take into account other sources of noise inherent to the imaging system such as read noise, which can be significant at high frame rates. Our analysis confirms that SNR is inversely related to frame rate and imaging frame rate should be carefully selected to optimize SNR in conjunction with the competing interest of maintaining high temporal resolution (Figure 7c). Furthermore, we identify the baseline fluorescence intensity of the nanosensor, F_0_, as an important parameter that influences SNR. SNR varies directly with F_0_ and nearly inversely with frame rate, with a local maximum at 10 Hz (Figure 7c). The observed maximum at 10 Hz arises from the diffusion induced broadening of a quantal release (Figure 2b, Figure 5a). For experiments wishing to optimize the temporal resolution of multiple dopamine firing events, as in Figure 5, choosing a nanosensor with strong baseline fluorescence will enable doing so via higher frame rates that achieve a decent fluorescent SNR.

The surprisingly strong dependence of SNR on F_0_ requires a closer examination. F_0_ depends on variables that are intrinsic to the fluorophore and to the imaging system as follows:
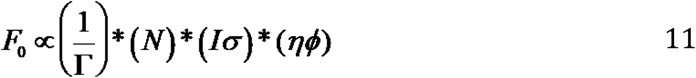

where Γ represents frame rate, N, the fluorophore number density, I, the excitation light source intensity and *σ*, the absorption cross-section of the fluorophore. The last two terms in equation 11 represent the quantum yield of the fluorophore (*η*) and the photon collection efficiency of the imaging system (*ϕ*). The direct dependence of SNR on F_0_ arises from noise filtering effects due to counting large numbers of photons, which inherently reduces Poisson noise. Thus, SNR optimization can be accomplished by tuning the fluorophore’s photophysical and chemical properties such as absorption cross-section, quantum yield, and analyte specificity. Two saturation regimes are worth noting: a neurotransmitter analyte saturation regime and a photon saturation regime. For the neurotransmitter concentration regime: the fluorophore number density, N, contributes to improved SNR only as long as the neurotransmitter analyte concentration does not become limiting. If the number of active binding sites on the nanosensor exceeds available analyte molecules, the relationship between N and SNR will deviate from that shown in Figure 7c. For the photon limiting regime: SNR will increase proportional to excitation intensity I, so long as photobleaching or fluorophore saturation does not dominate the imaging process. These latter effects demonstrate the importance of choosing optimal fluorophore excitation sources.

### Optimal Nanosensor Kinetic and Imaging Parameters can Record Behavior-Relevant Dopamine Dynamics for *In Vivo* Applications

The optical readout from nanosensors located in the striatal ESC will report on the space-averaged dopamine dynamics resulting from terminals in the volume surrounding the nanosensor. In practice, we wish to sample the cumulative behavior of dopamine over a region of interest in the ECS, for parameters relevant to our nanosensor. Our model fluorescent single-walled carbon nanotube sensors, with a 250 nm length and 1 nm width, diffuse through the ECS as rigid rods, and sample ECS subdomains on a short characteristic time scale of 200 ms.^52^ As such, the ensemble fluorescence modulation of optical nanosensors reflects average dopamine concentration. We simulate the ensemble fluorescence modulation of nanotube nanosensors in the ECS by averaging dopamine concentration over the simulation volume as described in Methods.

The volume averaged dopamine dynamics corresponding to 100 terminals firing phasically is shown in Figure 8a and Figure 8c. We define the phasic firing regime over a 2 second simulation with the physiologically-relevant spike train defined in Figure 4: A 4 Hz tonic firing rate for t = 0 to t = 0.4 s, a 20 Hz burst firing regime for t= 0.4 s to t = 0.7 s, a 0.5 s pause (0 Hz) until t = 1.2 s, and a 4 Hz tonic firing regime for 1.2 s to 2.0 s. Our simulation shows that synchronous firing of terminals (Figure 8d) yield transient dopamine concentrations in the ECS that range from 200 nM to 300 nM, with no basal levels between the peaks (Figure 8c, blue regions). Conversely, when neurons fire asynchronously (Figure 8b), tonic dopamine concentrations in the ECS fluctuate between 10 nM and 100 nM (Figure 8a, blue regions). The average tonic dopamine concentration in the striatal ECS obtained with our model is 50 nM (Figure S5), in line with results from prior computational studies^25^ and experimental measurements.^53,54^ This confirms that basal striatal dopamine is mediated by random, uncorrelated firing from dopaminergic terminals belonging to different neurons as opposed to correlated tonic firing. The volume averaging result is consistent with our simulations of many-terminal behavior presented in the previous section (Figure 4), validates our volume-averaging model, and corroborates previous experimental hypotheses about the nature of basal dopamine in the striatum originating from asynchronously firing neurons.

**Figure 8.**
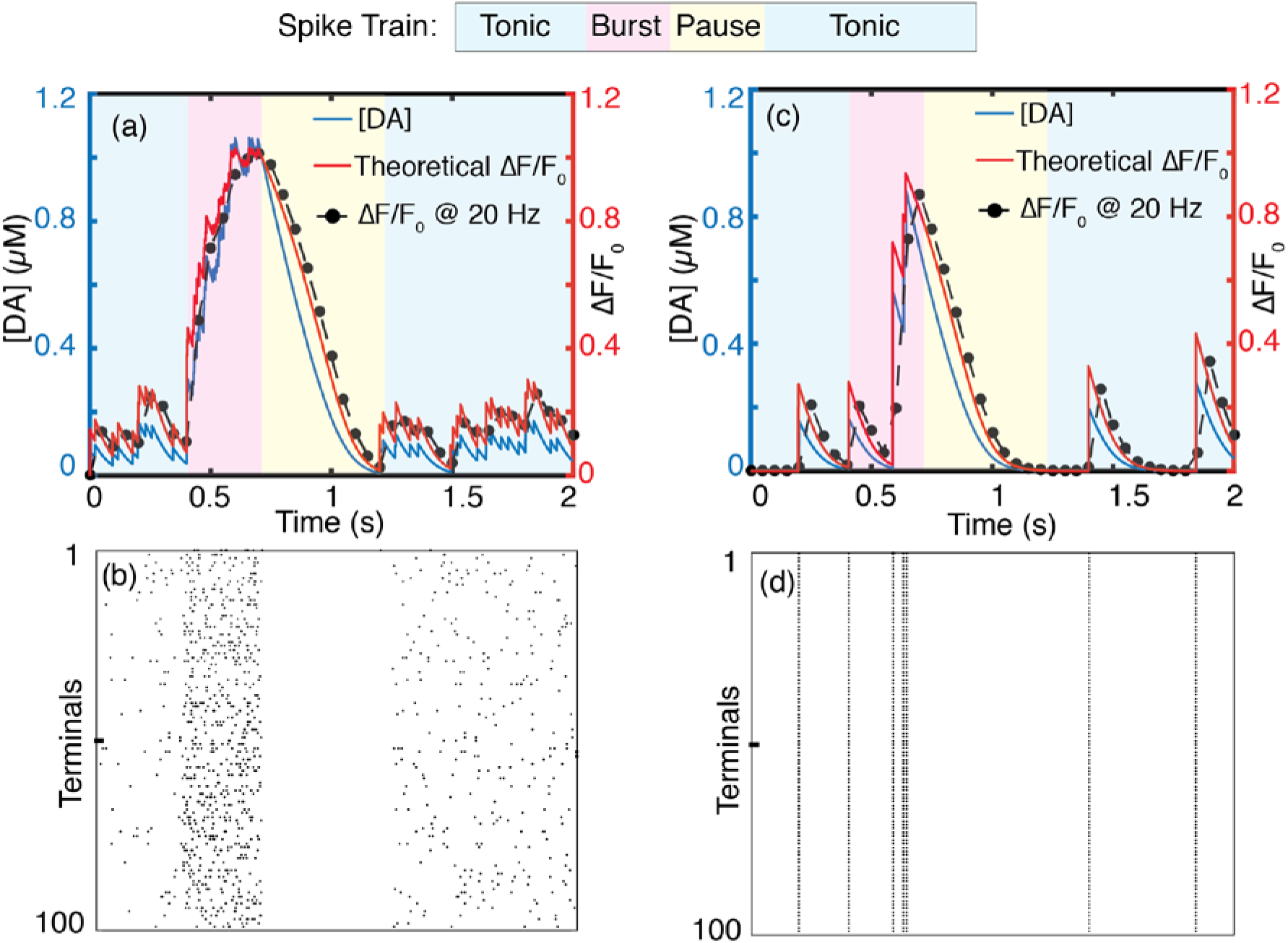
Volume averaged concentration profiles of 100 phasically firing dopamine terminals. (a) Dopamine concentration profile in which terminals fire asynchronously with theoretical and 20 Hz video-rate fluorescence nanosensor response profiles and corresponding (b) raster plot of synchronous firing activity corresponding to (a). (c) Dopamine concentration profile in which all terminals fire synchronously but phasically with theoretical and 20 Hz video-rate nanosensor responses and corresponding (d) raster plot showing the correlated activity of terminals in (c).

We next compute the ensemble dopamine nanosensor response profile for the theoretical versus practical cases of video-rate fluorescence imaging (Figure 8a, Figure 8c). We implement our results for a 20 Hz microscope frame rate identified previously as optimal for capturing striatal dopamine dynamics with optimal nanosensor parameters *K_eq_* = 1 (μM)^-1^, *α* = *2*. We present results for the two bounding cases of asynchronous firing (Figure 8a) and synchronous firing of all terminals (Figure 8c). During a firing burst that lasts 0.3 seconds, volume-averaged concentration rises to ~ 1 μM for both synchronous and asynchronous firing. These results represent space-averaged concentrations; locally, concentrations are heterogeneous and can be higher than the volumetric averages computed here (Figure 4a, b). Corroborating our prior results, imaging at 20 Hz, one can capture transient peaks during tonic firing in addition to the global concentration peak caused by a burst firing (Figure 8a, c). Furthermore, all behaviorally relevant firing regimes can be resolved, including the 0.5 s pause following the burst firing. At an imaging frame rate of 2 Hz, one can only resolve the concentration increase caused by burst firing; neither transient activity during tonic firing nor the pause following burst firing can be resolved (Figure S6). High affinity nanosensors saturate at tonic dopamine levels (Figure S7) whereas nanosensors with low dopamine affinity result in low nanosensor ΔF/F_0_ (Figure S8).

## CONCLUSIONS

The ECS constitutes an interconnected, porous and tortuous milieu that pervades neural tissue and serves as the medium through which neurons communicate with each other by way of chemical signaling. Our work quantifies the spatial and temporal nature of this chemical signal by using the dorsal striatum as a model system, and provides the requisite imaging and nanosensor kinetic parameters necessary to record chemical signaling in real time *in vivo*. Dopamine chemical signaling involves significant spillover of molecules from the synaptic cleft into the ECS, and a complex dynamic behavior arises as a consequence of release and simultaneous diffusion and reuptake. This work elucidates the dynamics by use of a rigorous, non-linear stochastic simulation, validated against existing experimental and computational literature. We show that the overflow of dopamine can be detected with optical probes placed in the ECS when proper imagining and kinetic parameters are chosen. A parameter space encompassing nanosensor kinetics and imaging frame rate is developed. Our work can be used to guide new nanosensor development, or to characterize those already developed. We use a generic receptor-ligand nanosensor kinetics that makes the results of our work broadly applicable to neurochemical imaging in the brain ECS. Furthermore, the model of the striatum developed here can be easily adapted to explore dynamics in other dopaminergic systems, such as the pre-frontal cortex and the nucleus accumbens, or to study analogous volume transmission phenomena of monoamines such as norepinephrine and serotonin. The simulation is modular and can be efficiently adapted to investigate broader variety phenomena that involve neurotransmitter dynamics such as pharmacokinetics of therapeutic agents and brain disease states.

## METHODS

### Discretization Scheme and Boundary Conditions

The model implements finite differences to solve the governing equation (equation 1). We take advantage of dopamine diffusion symmetry and isotropy to reduce the problem into 1D in spherical coordinates such that the distance from the release site, *r*, is the only spatial domain in the model. Symmetry at the site of a release site serves as a boundary condition for our numerical solution, and is used to calculate dopamine concentration at the center of the simulation volume. We provide details of the discretization scheme below.

The left hand side of the governing equation 1 can be written in difference form as:
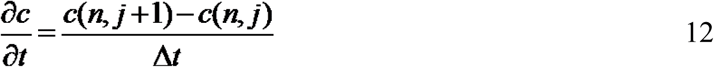

where indices *n* and *j* represent discrete steps in space and time, respectively.

To cast the right hand side of equation 1 in difference form, we first expand the Laplacian:
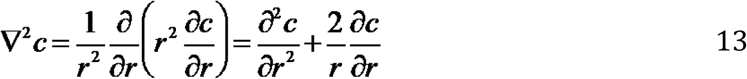

and discretize the first and second spatial derivatives as follows:
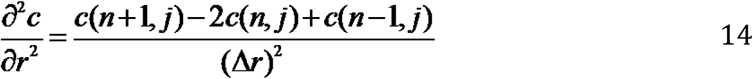

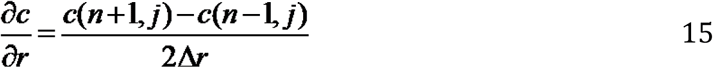

Using these difference equations and leaving the release and uptake terms as Q and U respectively, we can write an explicit equation for *c(n, j+1)* as (Equation 16):
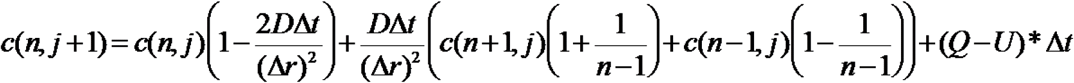

where *r*=*(n-1)Δr*.

The uptake term U is written explicitly in *c(n,j)* space as a Michaelis-Menten rate expression and the quantal release term (Q) is handled as described previously:
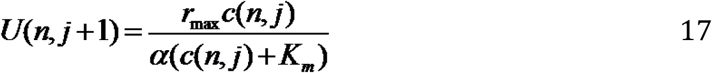

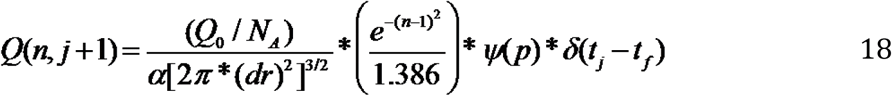

We discretize the symmetry boundary condition as follows. First we note that the governing equation as r → 0 becomes:
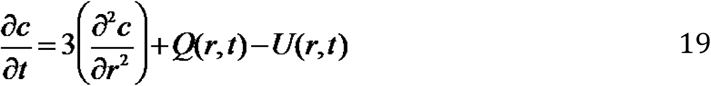

where we used L’Hopital’s rule to evaluate the limit. The spatial and temporal derivatives of equation 19 are discretized using equations 12 and 14 and then evaluated for n=1 (center), yielding:

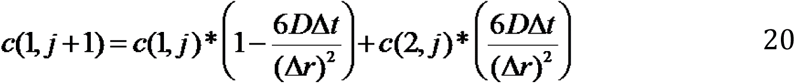

### Simulation Algorithm

At each simulation time step, the algorithm determines whether an action potential will invade a terminal based on a Poisson probability distribution with mean firing rate *F*. If there is a firing event, a quantal release of dopamine will occur based on a release probability, *p*, by toggling the binary variable *ψ*(*p*) between 1 (release) and 0 (no release). If a quantal release of dopamine occurs, dopamine concentration the volume immediately surrounding the terminal (*r= 0*) will be incremented by an amount in equation 3. Increment at subsequent volume elements are scaled as 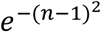, a factor that follows directly from a Gaussian probability density function. Once dopamine reuptake is determined, dopamine concentration at the location in space is decreased by an amount equal to the computed reuptake term. Because the discretized governing equation (eqn. 16) fails for *r=0 (n=1)*, where it becomes a singularity, we set Neumann’s symmetry principle at r=0 as a boundary condition and use it to compute concentration at the center (equation 20). We construct our simulation volume such that the effects of dopamine depletion at any point in space will result from dopamine reuptake by DATs within the simulation volume. Thus, we implement the Dirichlet boundary condition to enable modeling of dopamine reuptake effects at any distance from the center of the release site. Determining the exact location of this boundary requires solving the governing equation first, with a free boundary condition. We therefore set dopamine concentration to 0 when dopamine reuptake is higher than the available dopamine concentration at any given location from release point *r=0* as described by Berger et al.^55^ for the diffusion and uptake of oxygen in tissue. The simulation is implemented using MATLAB 2016a.

### Volume-Averaged Dopamine Dynamics

The dynamics from many terminals averaged over the volume encompassing the terminals is computed as follows:
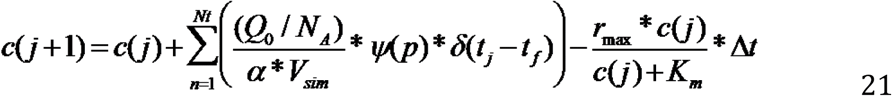

where c(j+1) is the volume averaged concentration of dopamine at time *t*_*j*+*1*_, c(j) is concentration at time *t_j_*, and *j* is the time index of the simulation. *Nt* is the number of terminals in the simulation volume *V_sim_* (1000 μm^3^). Terminals depolarize at designated times *t_f_*, where *t_f_* is the same for all terminals during synchronous firing and different for each terminal during asynchronous firing. The firing frequency sets the number of action potentials during a given simulation period. A Poisson distribution with a known mean firing rate sets the distribution of action potentials over the simulation time. Synchrony in firing activity is a reflection of the underlying functional connectivity of the ensemble. Synchronous firing (depolarization), however, does not mean all terminals release dopamine simultaneously; release of dopamine at each terminal is probabilistic and independent as per prior experimental literature,^15,56^ and thus set to 6% in our simulations. Note also that *N_t_* = *V_sim_ *ρ_t_* where *ρ_t_* is the density of dopamine terminals (Table 1) and *Δt* is the simulation time step (*Δt = t_j+1−_t_j_)*. The volume averaging as defined in equation 21 is valid only for large enough ensembles where diffusive flux out of the volume can be neglected. This is true if the volume is larger than the length scale of dopamine diffusion from a terminal. For small ensemble volume averaging, diffusive flux out of the volume needs to be taken into account because the length scale of the volume is smaller than the diffusion length scale of dopamine (Figure S5).

### Signal-to-Noise Ratio

The noise on the signal ΔF/F_0_ is related to the noise on F and F_0_. Using uncertainty propagation rules, we have:
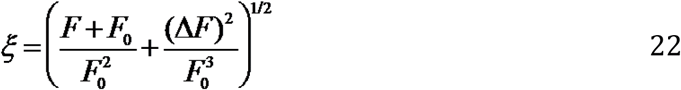

where *ξ* is the noise on our signal ΔF/F_0_. We use noise of 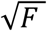 and 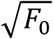 on F and F_0_ respectively, for Poisson limited imaging.

## SUPPORTING INFORMATION

**Figure S1** Peak dopamine concentration as a function of release site

**Figure S2** Non-linear uptake with *rmax* at 41 μM.s^-1^ compared to linear uptake with *rmax* at 4.1 μM.s^-1^

**Figure S3** Averaging N=20 simulation runs

**Figure S4** Dependence of SNR on the sensor parameter *K_eq_*.

**Figure S5** Volume-averaged behavior of N = 20 simulation runs

**Figure S6** Imaging at 2Hz fails to distinguish between behavior-relevant firing events of burst firing and pause in firing.

**Figure S7** Volume average dynamics imaged with a sensor with *K_eq_ = 100 μM^-1^*.

**Figure S8** Volume average dynamics imaged with a sensor with *K_eq_ = 0.1 μM^-1^*

## AUTHOR INFORMATION

**Corresponding Authors**

### Author Contributions

A.G.B. and M.P.L. conceived of the project and designed the study. A.G.B. developed the numerical analysis and model code. A.G.B and M.P.L developed the analysis of simulation results. I.R.M. and R.L.P. provided useful feedback during the course of model development. All authors discussed the results and commented on the manuscript.

### Notes

The authors declare no competing financial interest

## ACKNOWLEDGEMENTS

We thank members of the Wilbrecht, and Feller laboratories for insightful discussions. We thank the Molecular Graphics and Computation Facility at UC Berkeley College of Chemistry (NIH S10OD023532) for access to the computing facility during this work.

Notes and references

---

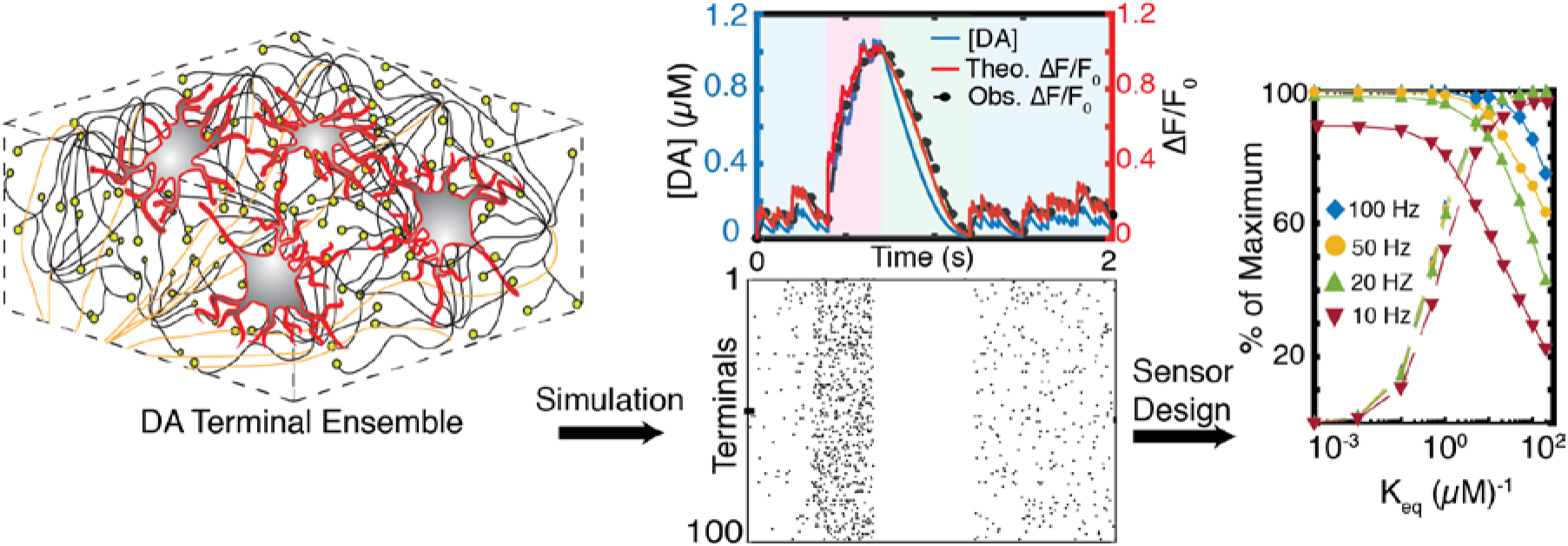
TOC GRAPHIC

